# Direct measures of liking and intensity of taste, smell, and chemesthetic stimuli are similar between young people reporting they did or did not have COVID-19

**DOI:** 10.1101/2023.10.07.561170

**Authors:** Emilia Leszkowicz, Katherine Bell, Amy Huang, Ha Nguyen, Danielle R. Reed

## Abstract

The recovery period from post-COVID-19 smell and taste dysfunctions varies substantially, lasting from a few days to over a year. We aimed to assess the impact of COVID-19 on post-COVID-19 chemosensory sensitivity in a group of young convalescents of eastern/central European ancestry. We measured subjects’ smell and taste capabilities with a standard testing kit, Monell Flavor Quiz (MFQ), and collected surveys on COVID-19 history. During testing, subjects rated liking and intensity of six odor samples (galaxolide, guaiacol, beta-ionone, trimethylamine, phenylethyl alcohol, 2-ethyl fenchol) and six taste samples (sucralose, sodium chloride, citric acid, phenylthiocarbamide, menthol, capsaicin) on a scale from 1 (dislike extremely, or no intensity) to 9 (like extremely, or extremely intense). There was no statistical difference in intensity ratings or liking of any sample between subjects who reported a history of COVID-19 (n = 34) and those reporting no history (n = 40), independent of presence/absence or severity of smell/taste impairments (*P* > 0.05). Additionally, neither vaccination status (full vaccination or no vaccination) nor time from the COVID-19 onset (2-27 months) correlated with liking or intensity. These results suggest that most young adults who had COVID-19 recovered their sense of smell and taste.

## Introduction

Chemosensory impairment accompanying COVID-19, which affects smell, taste, and chemesthesis, has been widely reported (Parma et al., 2020; Hannum et al., 2022). According to a 2020 meta-analysis, olfactory and gustatory dysfunctions affected around 44-53% of COVID-19 patients (Tong et al., 2020). Reports have shown that gustatory impairments are more frequent than olfactory ones, but rates vary depending on the country and region; for example, in New York, USA, 33% experienced anosmia, and 50% ageusia (Sehanobish et al., 2021); in California, USA, 68% anosmia, 71% ageusia (Yan et al., 2020); and in Brazil, 67% anosmia, 70% ageusia (Sampaio Rocha-Filho et al., 2022) (no data are available for Poland, the location of the current study). Ethnicity also appears to affect these rates (e.g., anosmia and ageusia in Caucasians, 55%; in Asians, 18%; von Bartheld et al., 2020), as does age (more prevalent dysfunctions in younger patients; Sehanobish et al., 2021) and date of infection onset (rates of dysfunction decreasing with later variants; Coelho et al., 2023).

A few mechanisms of smell dysfunction during COVID-19 have been suggested. They include, among others, SARS-CoV-2 infection of sustentacular cells and, to a lesser extent, Bowman’s glands, microvillar cells, and basal stem cells in the olfactory epithelium (Brann et al., 2020; Fodoulian et al., 2020); disrupted chromatin structure in olfactory sensory neurons (Zazhytska et al., 2022; Tan et al., 2023), and the release of pro-inflammatory cytokines (e.g., TNF-α) in response to viral infection leading to olfactory epithelium damage and dysfunction of olfactory sensory neurons (Torabi et al., 2020). In long-lasting smell dysfunctions, persistent inflammation, altered gene expression, and abnormalities of olfactory cleft and olfactory bulb have been observed, as well (Kandemirli et al., 2021; Finlay et al., 2022).

In turn, gustatory impairment has been associated with, inter alia, infection of salivary glands (Chen et al., 2020), epithelial cells on the tongue (Xu et al., 2020), and type II (but not type I or III) taste receptor cells in fungiform and circumvallate papillae; deficient stem/progenitor cell turnover in taste buds leading to impaired renewal of taste receptors cells (Doyle et al., 2021); or zinc insufficiency (Lozada-Nur et al., 2020). Inflammatory cytokines could also play a role; for example, interferons released in response to infection of tongue epithelial cells have been suggested to weaken regeneration of taste receptor cells (Wang et al., 2020).

A distinctive feature of COVID-19 anosmia and ageusia is its sudden onset and gradual improvement over a relatively short to moderate period of one to three weeks (Hornuss et al., 2020; Lee et al., 2020; Sampaio Rocha-Filho et al., 2022). However, anosmia and ageusia lasting more than one month or for three months (Kandemirli et al. 2021; Sampaio Rocha-Filho et al., 2022) and other smell/taste impairments lasting over one year (Boscolo-Rizzo et al., 2022) have also been reported. On the basis of statistical models and meta-analyses, an estimated 5-6% of patients with initial dysfunction might develop long-lasting persistent smell or taste dysfunction related to COVID-19 (Tan et al., 2022). Nonetheless, a recent paper by Nawab et al. (2024) with a predominantly (87%) white Hispanic and non-Hispanic population in the USA, suggests that chemosensory function after COVID-19 may fully recover.

The aim of our study was to assess whether COVID-19 left some lingering effects on smell and taste functions in a group of young convalescents of eastern/central European ancestry. We tested their smell and taste capabilities with a standard testing kit and compared their outcomes with those of a group reporting no history of COVID-19. We also collected surveys on COVID-19 history.

## Materials and methods

### Subjects

A total of 94 subjects (age, 21 ± 1 year; 17 males, 77 females) participated. The study followed the provisions of the Declaration of Helsinki for Medical Research Involving Human Subjects and was approved by the Bioethical Committee of the Regional Medical Chamber in Gdańsk, Poland. All subjects gave informed consent to participate in the study; they were not compensated for participation.

### Materials

The Monell Flavour Quiz (MFQ; Monell Chemical Senses Center, USA) contained six bottles of odorants labelled 1 to 6 (galaxolide, guaiacol, beta-ionone, trimethylamine, phenyl ethyl alcohol [PEA], and 2-ethyl fenchol), six bottles with samples of taste/chemesthetic stimuli labelled 7 to 12 (sucralose, sodium chloride, citric acid, phenylthiocarbamide [PTC], menthol, and capsaicin), and a paper response sheet. Samples concentrations and their characteristics are presented in Table 1. The odorants were present in beads in a volume of 0.2 mL/bottle (5 cellulose beads/bottle; Viscopearl, GPI Rengo, Japan), and the taste samples had a volume of 5 mL/bottle. The response sheet contained questions about liking of a sample on a scale of 1 (dislike extremely) to 9 (like extremely) and intensity of a sample on a scale of 1 (no intensity) to 9 (extremely intense), as well as what a sample smelled/tasted like (identity), which was not included in our analyses.

**Table 1.**
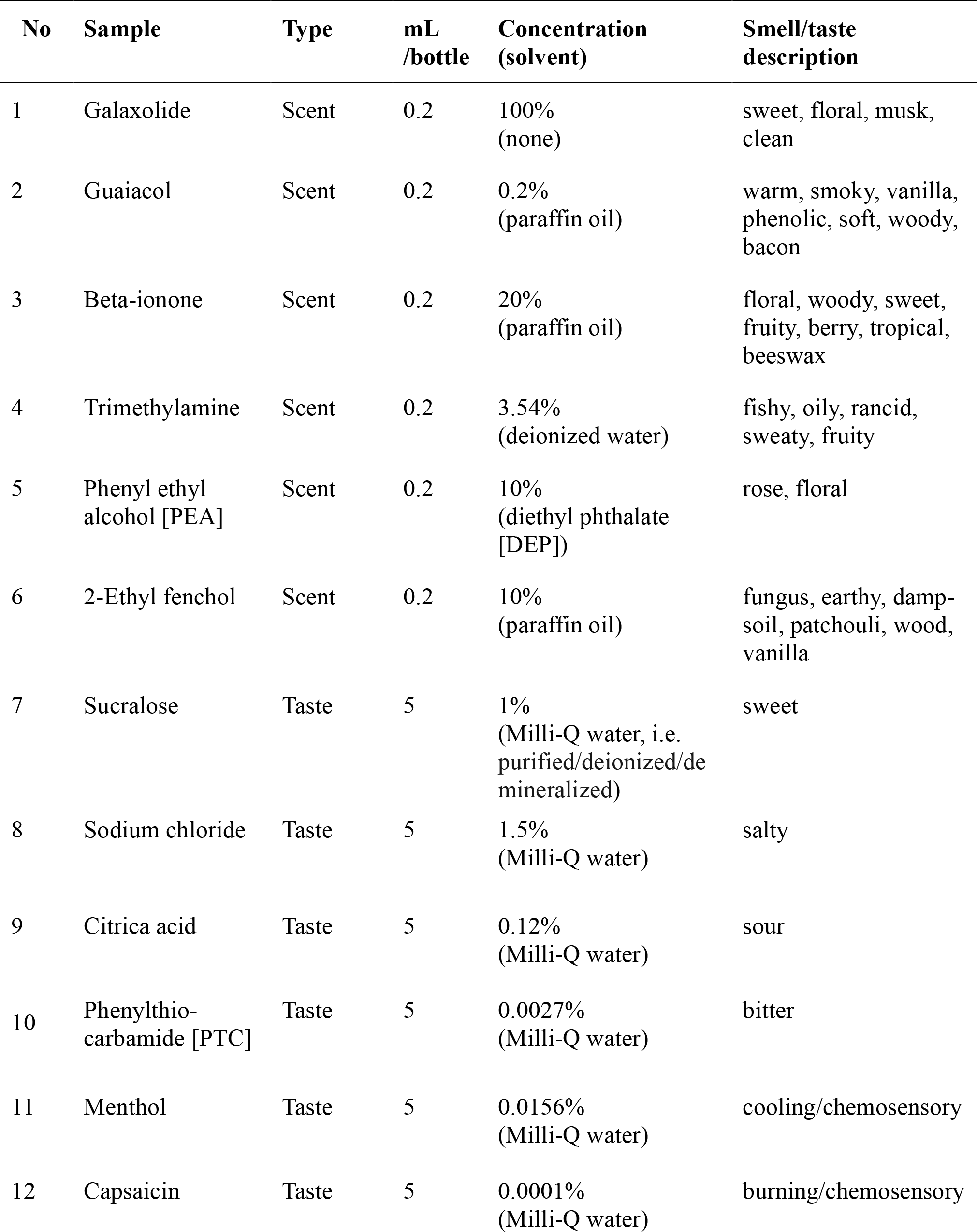
The concentrations of samples used in the Monell Flavour Quiz, and their characteristics.

| No | Sample | Type | mL<br>/bottle | Concentration<br>(solvent) | Smell/taste<br>description |
| --- | --- | --- | --- | --- | --- |
| 1 | Galaxolide | Scent | 0.2 | 100%<br>(none) | sweet, floral, musk,<br>clean |
| 2 | Guaiacol | Scent | 0.2 | 0.2%<br>(paraffin oil) | warm, smoky, vanilla,<br>phenolic, soft, woody,<br>bacon |
| 3 | Beta-ionone | Scent | 0.2 | 20%<br>(paraffin oil) | floral, woody, sweet,<br>fruity, berry, tropical,<br>beeswax |
| 4 | Trimethylamine | Scent | 0.2 | 3.54%<br>(deionized water) | fishy, oily, rancid,<br>sweaty, fruity |
| 5 | Phenyl ethyl<br>alcohol [PEA] | Scent | 0.2 | 10%<br>(diethyl phthalate<br>[DEP]) | rose, floral |
| 6 | 2-Ethyl fenchol | Scent | 0.2 | 10%<br>(paraffin oil) | fungus, earthy, damp-<br>soil, patchouli, wood,<br>vanilla |
| 7 | Sucralose | Taste | 5 | 1%<br>(Milli-Q water, i.e.<br>purified/deionized/de<br>mineralized) | sweet |
| 8 | Sodium chloride | Taste | 5 | 1.5%<br>(Milli-Q water) | salty |
| 9 | Citric acid | Taste | 5 | 0.12%<br>(Milli-Q water) | sour |
| 10 | Phenylthio-<br>carbamide [PTC] | Taste | 5 | 0.0027%<br>(Milli-Q water) | bitter |
| 11 | Menthol | Taste | 5 | 0.0156%<br>(Milli-Q water) | cooling/chemosensory |
| 12 | Capsaicin | Taste | 5 | 0.0001%<br>(Milli-Q water) | burning/chemosensory |

Non-carbonated spring water containing no taste- and no odor-defining substances (Cisowianka, Nałęczów Zdrój, Poland) and an unsalted wheat matzo (Sante, Warsaw, Poland) were provided to cleanse the palate between taste samplings. The questionnaire included questions about sex, ethnicity, race, and COVID-19 history such as COVID-19 symptoms, duration, and timing.

### Procedure

Subjects were asked to refrain from eating, drinking (apart from water), and smoking for at least one hour before sensory testing. They filled out the COVID-19 questionnaire online after the sensory testing.

For sensory testing, subjects were instructed to rate the smell or taste of samples 1-12 and to mark their responses for liking, intensity, and identity for each sample on the response sheet. For samples 1-6, they were instructed to hold a bottle 2.5 cm (1 inch) from the nose and sniff several times. For samples 7-12, they were asked to rinse their mouth with water and spit the water out; then, to close their noses with a nose clip, pour the entire amount of each sample into the mouth, hold it in the mouth for approximately five seconds, and spit out the solution into an empty cup. Between each taste sample, they rinsed their mouth with water and ate a matzo to remove any lingering taste from the previous sample.

### Data analyses

Our goal was to classify subjects into two groups: those who had COVID-19 one or more times and those who never had COVID. Some people were difficult to classify, so we first applied the following exclusion criteria: (1) a negative SARS-Cov-2 test or no such test, and some disease symptoms but no smell/taste disturbances, (2) inconsistent responses or not following instructions in the COVID questionnaire, or (3) not sure if they had COVID-19 or “other” in a response to a question about COVID-19. After applying the exclusion criteria, 74 subjects were included in data analyses (14 males, 60 females), and they were divided into two groups based on the questionnaire responses: relaxed-COVID group, who had a positive SARS-Cov-2 test, or either had a negative SARS-Cov-2 test or did not take the test but reported they had had COVID-19 with accompanying smell/taste impairment (n = 34); and no-COVID group, who reported they had not had COVID-19 (no SARS-Cov-2 test, and no symptoms; n = 40). Additionally, to reflect the degree of certainty for the COVID-19 diagnosis, we created a strict-COVID subgroup within relaxed-COVID group, comprising only those who had a positive SARS-Cov-2 test (n = 16). It should be noted that during the early stage of the pandemic, SARS-Cov-2 tests were unavailable or sparse, therefore it was not possible to require a confirmatory test for early COVID-19 infections.

To assess the influence of COVID-19 on liking and intensity of samples, we compared strict-COVID and relaxed-COVID groups with no-COVID group. Additionally, in the relaxed-COVID group, we analyzed if liking and intensity were affected by (a) vaccination status during COVID-19 (full vaccination. with at least two COVID-19 vaccinations, or no vaccination; n = 13 and n = 19, respectively), (b) chemosensory disorders, that is smell disorder status during COVID-19 (total anosmia, other smell disturbances, no smell disturbances; n = 19, n = 8, n = 7, respectively), and taste disorder status during COVID-19 (total ageusia, other taste disturbances, no taste disturbances; n = 18, n = 7, n = 9, respectively), and (c) time since COVID-19 (in months).

A Shapiro-Wilk test was used to test data normality distribution for choosing an appropriate statistical method. The distribution of most variables (apart from citric acid intensity, W = 0.957, *P* = 0.012) departed significantly from normality (from W = 0.952, *P* = 0.007 for guaiacol intensity to W = 0.711, *P* < 0.001 for capsaicin intensity). Based on this outcome, we used Mann-Whitney U and Kruskal-Wallis non-parametric tests for between-group analyses and used medians with interquartile range to summarize variables. Correlations were calculated as a Spearman rank *R* coefficient.

To decide which multiple testing correction to apply, we looked for and found no clear correlations among the outcome measures. To avoid overly stringent correction of multiple analyses, false discovery rate (FDR) correction (with FDR < 5%) was applied to correct for multiple comparisons in between-group analyses, with the original α < 0.05 and the total number of comparisons 24 (liking and intensity of 12 samples).

To evaluate the statistical power of the tests in a clear and informative, albeit approximate, way, the average Cohen’s *d* effect size and the average test power were used to draw accurate conclusions about differences between the relaxed-COVID and no-COVID groups. The average Cohen’s *d* effect size for the 12 stimuli was calculated on modules of individual stimulus effect sizes. Test power was computed with a statistical power calculator (Power T Z Calculator; statskingdom.com).

## Results

All subjects included in analyses were second-year students of the same age (median, 21 years; range, 20-22 years; 81% women; 100% eastern/central European ancestry).

### COVID-related taste and smell disorders

The disorders in the relaxed-COVID group are presented in Table 2. Among 16 subjects in the strict-COVID group (with a positive SARS-Cov-2 test), 7 (44%) did not have accompanying smell or taste disorders: one of them was diagnosed with COVID-19 in early 2000, two in late 2021, and four in early 2022.

**Table 2.** Number of participants who reported smell and/or taste disorders in the relaxed-COVID group (n = 34, percent of ‘n’ in brackets), and their self-reported smell and/or taste recover time (n = 27, percent of ‘n’ in brackets).

|  |  |
| --- | --- |
| Smell & taste impairments | 27 (79%) |
| Smell only impairments (taste fine) | 2 (6%) |
| Taste only impairments (smell fine) | 0 |
| Total anosmia | 19 (56%) |
| Total ageusia | 18 (53%) |
| 2-week recover time | 17 (63%) |
| (1-3)-month recover time | 5 (18.5%) |
| Smell/taste still disturbed after $\geq 6$ months | 5 (18.5%) |

Self-reported recovery time from smell and/or taste disorders in the relaxed-COVID group is shown in Table 2. Out of five subjects who did not regain their sense of smell or taste after 6 months or longer, on the day of sensory testing four reported poorer smell and taste sensitivity than before COVID-19 (time post-COVID-19 onset: 6, 13.5, 19, and 19 months) and one reported only weaker smell sensitivity (19 months post-COVID-19 onset). These five were not differ in terms of samples’ preference or intensity ratings from those who did not report such lingering post-COVID effects. Three other subjects reported that some products that were previously fine now smelled/tasted bad after COVID-19 (4, 7, and 21 months post-COVID-19 onset), and one subject reported liking what used to be disgusting to them (18 months post-COVID-19 onset). Smell impairments apart from total anosmia included phantosmia, hyposmia, smell fluctuations (appearing/disappearing), and selective loss of smell. Taste impairments besides total ageusia included selective changes/loss in taste modalities, and hypogeusia.

### COVID vs no-COVID

For ratings of liking and intensity of the samples, no difference was found between strict-COVID and no-COVID groups (data not shown) or between relaxed-COVID and no-COVID groups (Fig. 1), after FDR correction. Liking and intensity for relaxed-COVID vs no-COVID: U = (499 to 676), Z score with continuity correction = (−2.010 to 1.102), *P* = (0.044 to 0.969 before FDR correction). Thus, because neither strict-COVID nor relaxed-COVID groups differ from no-COVID group, we limited our further within-group analyses reported below to the relaxed-COVID group.

**Figure 1.**
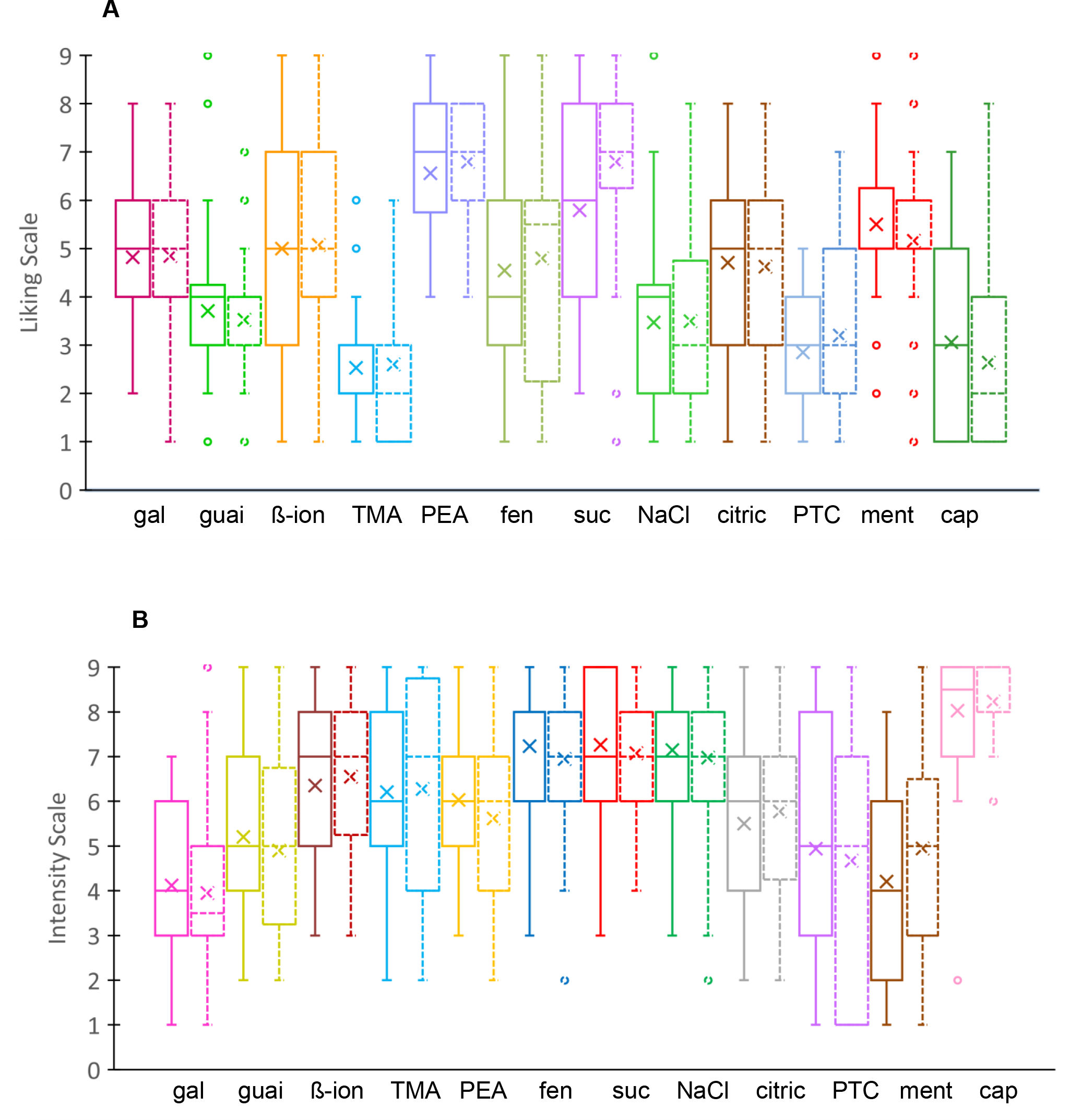
Liking (A) and intensity (B) for samples tested by subjects who reported having COVID-19 (relaxed-COVID group; solid lines) and those who did not (no-COVID group; dashed lines). Abbreviations: gal, galaxolide; guai, guaiacol; ß-ion, beta-ionone; TMA, trimethylamine; PEA, phenyl ethyl alcohol; fen, 2-ethyl fenchol; suc, sucralose; NaCl, sodium chloride; citric, citric acid; PTC, phenylthiocarbamide; ment, menthol; cap, capsaicin. In these box plots, the rectangle indicates the interquartile range, the middle bar the median, the x the mean, and the whiskers the range. For all 12 samples *P* > 0.05, Mann-Whitney U test with FDR correction for multiple comparisons.

### Vaccination status

In the relaxed-COVID group, no difference was found between subjects who were fully vaccinated during COVID-19 and those who were not vaccinated, after FDR correction. Liking or intensity of samples did not differ by vaccination status: U = (68 to 123), Z score with continuity correction = (−2.171 to 1.777), *P* = (0.030 to 1.000 before FDR correction).

### Chemosensory disorders and time since COVID-19

The status of smell disorder or taste disorder during COVID-19 had no effect on liking or intensity of the samples in the relaxed-COVID group (*P* > 0.05). During the sensory testing, subjects were 1 to 29 months post-COVID-19. No correlation was found between time since COVID and subjects’ ratings of sample liking or intensity (Spearman rank R coefficient = -0.243 to 0.323).

### Effect size and power calculations

The average of Cohen’s *d* effect sizes for all stimuli for relaxed-COVID vs no-COVID was 0.150 (0.145 for liking, 0.155 for intensity), and the average test power size was 0.097 (0.094 for liking, 0.101 for intensity). Based on these outcomes, a sample size of 1,380 subjects per group would be needed to detect statistically significant between-group differences, considering the small average effect size and test power size, and assuming a significance level of α < 0.002 after Bonferroni correction for multiple comparisons and a test power of 0.8 (using Power T Z Calculator).

## Discussion

Our results show that young people of eastern/central European ancestry who reported previous COVID-19 infection with or without accompanying chemosensory dysfunctions did not differ in liking or intensity ratings of smell, taste, or chemesthetic samples from those who reported not having COVID-19. Time since COVID-19 onset also did not correlate with liking or intensity of samples. Similarly, neither vaccination status nor severity of smell/taste disturbances related to COVID-19 had an effect on those measures.

Most convalescents reported they had regained their sense of smell and taste within two weeks after COVID-19 onset; others had smell/taste impairments for one to three months. Interestingly, self-reports on current smell/taste/chemesthetic sensitivity did not always match measures obtained with the MFQ: for a few subjects who reported that 6-19 months post-COVID-19 they still had a poorer sense of smell or of smell and taste than before infection, ratings of samples intensity or liking did not differ from others who did not report such deficits. Even though this suggests that those subjects had some better smell and/or taste capabilities before COVID-19, lack of data on their pre-COVID-19 chemesthetic capabilities makes it impossible to precisely define to what degree their deficits were COVID-related, and what smell/taste qualities, and quantities were affected by COVID-19. This result also argues for the need for regular smell and taste screenings raised in other reports (Mainland et al., 2020; Hunter et al., 2023). Development and widespread use of relevant tests can benefit society, especially that it might help in early diagnosis of diseases in which smell loss is one of the symptoms, such as neurodegenerative diseases and accelerated cognitive decline (Albers et al., 2006; Dintica et al., 2019; Doty and Hawkes, 2019), or head trauma (Callahan and Hinkebein, 1999).

Potential differences in smell and taste capabilities between COVID-19 convalescents and those who did not have COVID-19 were also explored in a recent paper by Nawab et al. (2024). Their study was performed in mainly white population (87%) of Hispanic and non-Hispanic ethnicity, residing in the USA; participants were older than in our study, that is mean age in their experimental and control groups were 42 (95 % CI: 36–47), and 46 (95 % CI: 39–52), respectively; and sensory testing was done with UPSIT test (smell), and the Burghart Taste Strips Test (taste). It is worth noting that despite these methodological differences between our and Nawab’s et al. studies, our results are congruent with their results, which were the full recovery of chemosensory functions in post-COVID patients.

Olfactory sensory neurons and receptor cells in taste buds regenerate in mammals (rodents) on average in 30-60 and 8-12 days, respectively (Beidler and Smallman, 1965; Mackay-Sim and Kittel, 1991). Experiments on hamsters revealed that the olfactory epithelium could recover from SARS-CoV-2 infection in 10-21 days, depending on the epithelial region (Urata et al., 2021). The turnover rate of taste receptor cells and recovery period of the olfactory epithelium could partially explain why anosmia, ageusia, and other smell/taste disturbances were fully reversed in most subjects in two weeks. On the other hand, impairments of basal stem cells in olfactory epithelium and stem/progenitor cells of taste buds, olfactory cleft inflammation, olfactory bulb impairments (including its degeneration), to name a few, could account for a longer renewal process for other subjects in this study (Kandemirli et al., 2021; Doty, 2022; Finlay et al., 2022).

We found no clear difference in the liking or intensity rating of the samples between subjects who self-reported lasting poorer sense of smell/taste/chemesthesis and the others, which suggests that weaker chemesthetic sensitivity does not necessary apply to all stimuli and seems to be limited to some of the stimuli encountered by subjects in everyday life. That in turn may reflect different recovery times for diverse modalities of smell, taste, or chemesthesis.

Taste receptor cells, which discriminate various taste modalities, differ in life-spans: type II cells, which discriminate sweet, bitter, and umami, have a life-span of about 8 days; type III cells, which discriminate sour, 22 days; and type I (glial-like) cells, 8-24 days; the life-span of salty-discriminating cells, which do not meet the criteria of any of the three types, remains unknown (Perea-Martinez et al., 2013; Nomura et al., 2020). Similarly, the regeneration rate of the olfactory epithelium varies depending on its zone: the epithelium in the regions of the nasal septum, and medial turbinate recovers fastest (in 21 day; hamsters), and in dorsal and lateral turbinates more slowly (not fully recovered after 21 days) (Urata et al., 2021). Moreover, vulnerability to viral infection varies in these regions, with the medial turbinate being least vulnerable (Urata et al., 2021). Heterogeneous life-spans of taste receptor cells and nonuniform regeneration rates of different olfactory epithelium zones could be one of the reasons that various taste or smell qualities and intensities do not always regenerate at the same time. Hence, the self-reported generally weakened sense of taste/smell may not necessary be mirrored in the intensity rating of a particular sample. Additionally, it is worth noting that homeostasis of taste bud declines in aging (see Feng et al., 2014), and our subjects were young, which could partially account for their recovery rate.

COVID-19-related smell and taste recovery time may be longer than 6 months (Hopkins et al., 2021; Nguyen et al., 2021), and even one year post-infection smell and taste impairments have been observed (Boscolo-Rizzo et al., 2021). That is not surprising because some post-viral olfactory dysfunctions are known to recover after more than 24 months (Lee at al., 2014).

Our post-COVID-19 subjects who had been affected with smell and taste dysfunctions did not differ in liking or intensity rating of tested samples from controls who had not have COVID-19. That finding, together with the fact that subjects who were at least 24 months post-COVID-19 (n = 3) did not report any lingering effects of smell or taste disturbances, suggests that chemosensory impairments linked to SARS-CoV-2 infection may be fully reversible, especially in younger people, but the recovery process may take a long time.

## Acknowledgments

We thank students from the Faculty of Biology at the University of Gdańsk, especially Joanna Pasztelan, for their assistance in data collection in Gdańsk (Poland).

## Data availability

The data underlying this article will be shared on reasonable request to the corresponding author.

